# Disrupted endothelial cell–cell dynamics in ALK1 and SMAD4 deficiency drive arteriovenous malformations

**DOI:** 10.1101/2025.11.26.690643

**Authors:** David Hardman, Olya Oppenheim, Wolfgang Giese, Holger Gerhardt, Miguel O. Bernabeu

## Abstract

Arteriovenous malformations (AVMs) are a hallmark of hereditary haemorrhagic telangiectasia (HHT) and arise from abnormal vascular remodelling. Although AVM formation has been associated with disruptions in the BMP9/10 signalling pathway, the distinct contributions of its components remain unclear. Here, we combine *in vitro* mosaic endothelial cell (EC) cultures with agent-based modelling (ABM) to investigate how knockdown of the BMP9/10 pathway components ALK1 and SMAD4 alters cell-cell interactions and collective vascular organisation.

Using cell tracking data, we inferred the mechanical interactions between neighbouring ECs in 2D monolayers by applying Approximate Bayesian Computation to an ABM of cell migration. SMAD4 knockdown increased the motile forces generated by ECs, while ALK1 knockdown weakened the ability of cells to push apart and rearrange with their neighbours, both resulting in greater mixing and fluidity within the cell layer. When these altered interaction profiles were incorporated into an ABM of collective EC dynamics during vascular remodelling, they gave rise to distinct AVM-prone phenotypes. SMAD4-deficient populations exhibited local blockages as highly polarised cells migrated excessively, whereas ALK1-deficient populations disrupted vascular patterning by impairing coordinated polarity and neighbour separation, favouring flow reversal.

Our findings suggest that canonical BMP9/10 pathway defects destabilise the biomechanical balance of endothelial interactions, leading to emergent collective behaviours that predispose vessels to AVM formation.

**Significance Statement:** Arteriovenous malformations (AVMs) are abnormal connections between arteries and veins that disrupt blood flow and can cause stroke or life-threatening bleeding. They are a hallmark of hereditary haemorrhagic telangiectasia (HHT), a genetic vascular disorder, but the cellular events that initiate AVMs remain unclear. Using a combination of endothelial cell cultures and computational modelling, we show that loss of two key HHT-linked genes, ALK1 and SMAD4, disrupts the coordinated movement and interactions of endothelial cells. Although both deficiencies lead to AVM-like behaviours, they do so through distinct mechanisms. SMAD4 loss drives excessive collective migration, whereas ALK1 loss impairs polarity and neighbour separation. These findings provide a mechanistic framework for how genetic defects could contribute to cell-cell dynamics observed in vascular malformations.

## Introduction

Vascular remodelling is essential for shaping functional blood vessels, ensuring efficient haemodynamics and tissue perfusion. When remodelling is disrupted, abnormal vessel connections known as arteriovenous malformations (AVMs) can arise, with serious clinical consequences. Blood flow^1^ and endothelial cell (EC) behaviour play a significant role in this remodelling process, polarising and migrating against the direction flow in response to vessel wall shear stress (WSS)^2^.

AVMs are a hallmark of hereditary haemorrhagic telangiectasia (HHT), an inherited vascular disorder. Type 2 HHT is caused by loss-of-function mutations in the endothelial surface receptor Activin-receptor-like kinase 1 (*AVCRL1* encoding ALK1)^3^. SMAD4 functions downstream of ALK1 in the Bone Morphogenic Protein (BMP) 9/10 pathway and of ALK5 in the Transforming Growth Factor Beta (TGF-β) pathway, and is a shared mediator in activin signaling. Mutations in the *SMAD4* gene are present in families with Juvenile Polyposis/HHT syndrome. Deletion of both endothelial *Alk1* and *Smad4* in mouse models^4^ have resulted in AVM formation and lethality though the relative effects of ALK1 and SMAD4 on AVM formation and the mechanisms by which they mediate EC behaviour in AVM formation remain poorly understood. While these animal models have been invaluable, they do not allow direct live imaging of AVM initiation within intact vasculature, meaning that the cell–cell dynamics underlying AVM formation have not yet been described in detail in vivo.

To determine the influence that inhibiting ALK1 and SMAD4 has on healthy cells, we conducted *in vitro* experiments with human umbilical venous endothelial cells (HUVEC) on mosaic populations combining knockdowns of either ALK1 or SMAD4 with control cells and tracked the dynamics of cell motion over time.

Previous work by Edgar *et al*^5^ applied an agent-based model (ABM) of EC motion and mechanical cell–cell interactions on idealised vascular networks to investigate the collective dynamics of cells during vascular remodelling. Distinct regimes of EC force dynamics favourable to AVM creation were identified based upon theoretical forces of cell motility, contractility, and adhesion, together with the proportion of cells moving with or against the direction of WSS. A similar modelling approach based on self-propelled particles (SPP) was used to study remodelling behaviour in the yolk sac vasculature of quail embryos and compared directly with microscopic images, demonstrating the predictive power of such modelling approaches^6^.

Here, we used cell tracking data from our *in vitro* mosaic experiments with the level of granularity needed to parameterise an ABM of endothelial interactions in 2D monolayers for the first time. By systematically varying the range of cell–cell adhesion (“cohesion”), neighbour-displacing forces that separate adjacent cells (“extrusion”), and directed motility (“migration”) in the ABM we can use Approximate Bayesian Computation with Sequential Monte Carlo sampling (ABC-SMC) to infer the likely distribution of interaction forces associated with cell adhesion, contractility, and motility to recapitulate the experimental observations. This parameter inference method is applied to EC populations with knockdowns of ALK1, SMAD4 and a control to determine the difference in cell mechanical interaction regimes between the different populations. This will provide insight into forces at cell-cell junctions, which are important for maintaining vascular integrity^7^.

We found that SMAD4 knockdown cells exhibited increased motile force generation, whereas ALK1 knockdown weakened the forces separating neighbouring cells that normally allow cell rearrangement within the monolayer. Together, these changes reduced the efficiency of neighbour displacement, making it easier for cells to intermingle without the normal checks on rearrangement.

Having investigated force transmission in 2D cell monolayers and characterised the impact that ALK1 and SMAD4 have on them, we next ask the question of whether these changes can mechanistically explain AVM formation. Our ABM model of EC collective dynamics during vascular remodelling was parameterised with the experimentally derived proportional changes in cell motility, adhesion, and neighbour displacement forces observed between control, ALK1 knockdown and SMAD4 knockdown cells. Loss of flow and reversal of flow in remodelling networks have previously been postulated to be associated with shunt formation^5^. By quantifying the proportion of vessels in a network in which flow is lost or reversed over time we can determine whether the changes in cell adhesion and directional migration associated with each knockdown are predictive of AVM formation. Our simulations indeed predict that AVM are more likely to arise from the loss of either SMAD4 or ALK1, through distinct mechanisms.

Taken together, our combined experimental-computational approach describes changes in EC force dynamics in SMAD4 and ALK1 deficient cells and predicts that deficiencies of either will create emergent conditions favourable to AVM formation *in vivo*.

## Results

### Cell motion and intercellular force coefficients from *in vitro* EC mosaic experiments

Control and ALK1 knockdown cells were measured to be of similar sizes (cell width 15μm) while SMAD4 knockdown cells were larger (cell width 19μm). Measurements of cell direction of motion with respect to flow and inference of intercellular force coefficients were obtained from live images from *in vitro* mosaic experiments of control/control and control/knockdown EC monolayers exposed to flow for 36 hours (Fig. 1A).

**Figure 1.**
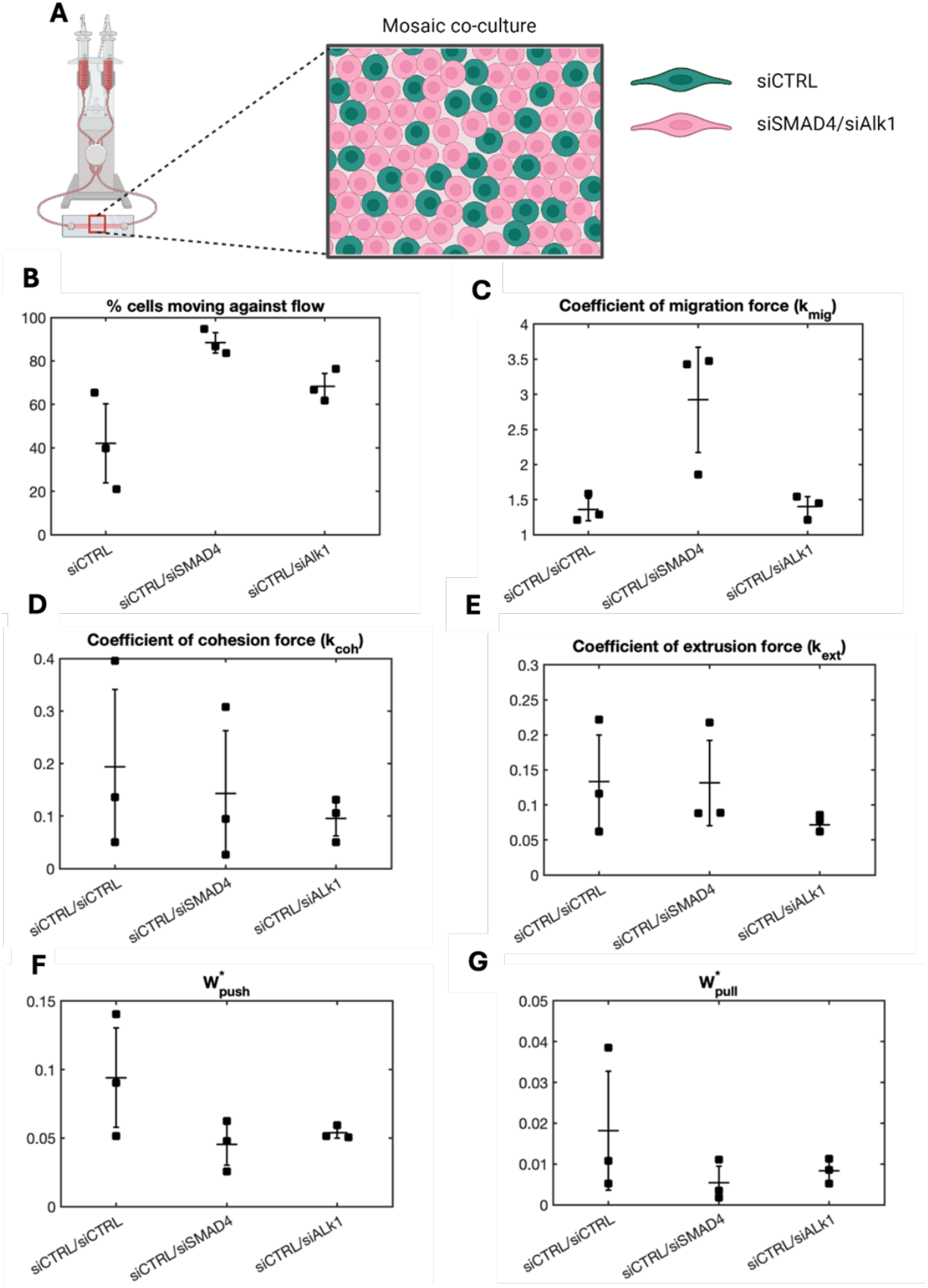
*Figure 1:* Measurements of cell direction of motion and inferred intercellular force. **(A)** Sketch of fluid pump system (left) and illustration of stained endothelial cell mosaic, siCTRL in green and siALK1 cells in magenta (right). (**B**) Percentage of cells moving against the direction of flow (**C-E**) Inferred coefficients of intercellular forces. (**F**) Work required to push neighbouring cells together (W ^*^_push_). (**G**) Work required to pull neighbouring cells together (W ^*^_pull_). Data for plots B-G obtained from cells after 36 hours of exposure to 6 dyne/cm^2^ unidirectional flow (mean +/-1 standard deviation)

For cell mosaic experiments with control/SMAD4 knockdown, an average of 88.3 +/-4.7% cells were moving against the direction of flow, a greater proportion than the control/ALK1 knockdown experiments (68.2 +/-6.1%) (Fig.1B). In control/control mosaic experiments an average of 42.0 +/-18.2% moved against flow, though this proportion was found to rise as the cells were exposed to flow beyond 36 hours (see Supplementary materials, Table S1).

The coefficient of cell migration force (*k*_*mig*_) in the control/control and control/ALK1 knockdown mosaic experiments was inferred to be similar in magnitude (1.35+/-0.16 and 1.40+/-0.14 respectively) while *k*_*mig*_ in the control/SMAD4 knockdown cells was found to be higher (2.92+/-0.75) (Fig. 1C). The average coefficient of cell cohesion force (*k*_*coh*_) was slightly lower in the control/ALK1 knockdown cells (0.10+/-0.03) compared to control/SMAD4 and control/control cells (0.14+/-0.12 and 0.19+/-0.15 respectively) (Fig. 1D). The large ranges of *k*_*coh*_ in control/SMAD4 knockdown and control/control cells suggest that any differences in cohesion force are not significant. Control/ALK1 knockdown cells showed cell extrusion force (*k*_*ext*_) to be consistently low (0.08+/-0.01) compared to control/control and control/SMAD4 knockdown cells (0.13+/-0.07 and 0.13+/-0.06 respectively).

From equation (1), the average work required to push two neighbouring cells apart W^*^_push_ (Fig. 1E) was found to be lower for both control/SMAD4 knockdown (0.045+/-0.015) and control/ALK1(0.054+/-0.004) knockdown cells than the control/control cell experiments (0.094+/-0.036). In the case of control/SMAD4 knockdown this is due to the high *k*_*mig*_ while in control/ALK1 knockdown this is due to the lower *k*_*ext*_. Assuming cell yield stretch is constant, the average work required to pull two neighbouring cells together W^*^_pull_ was found to be slightly lower in control/SMAD4 knockdown (0.005+/-0.004) and control/ALK1 knockdown (0.008+/-0.003) than control/control cells (0.018+/-0.015) (Fig 1G).

### Experimental force measurements predict AVM-like behaviours in SMAD4 and ALK1 knockdowns

Previous work^5^ quantifying EC behaviour in ‘honeycomb’ networks of vessels simulating vascular remodelling found that AVM formations were associated with increases in the proportion of vessels with flow loss and flow reversal. In these models, flow disturbances arose due to the accumulation of ECs which blocked or diverted blood flow.

It was also shown that varying the magnitude of W^*^_pull_ affects the proportion of vessels with both flow loss and flow reversal in a manner dependent upon the proportion of cells moving against the direction of blood flow. Here, the experimentally derived intercellular force parameters (see Table 1) and cell direction of motion with respect to flow from *in vitro* experiments were applied to simulated honeycomb vessel networks and changes in vessels with flow drop out and flow reversal were recorded. Proportions of cells moving with or against flow changed significantly over time for control only cells populations in *in vitro* experiments (see Supplementary Materials Table S1). Values of proportional cell direction were therefore taken from results after 60 hours in flow, rather than the 36 hours recorded above, to allow cell motion to reach an equilibrium.

**Table 1.** Intercellular force coefficients inferred from in vitro experiments and converted to vascular networks using equation (4).

| Experiment | Force coefficients from *in vitro* measurement |  |  | Force coefficients for Vascular networks |  |  |
| --- | --- | --- | --- | --- | --- | --- |
| | $k_{mig}$ | $k_{coh}$ | $k_{ext}$ | $k_{mig}$ | $k_{coh}$ | $k_{ext}$ |
| siCTRL/siCTRL | 1.36 | 0.194 | 0.133 | 3 | 0.333 | 3 |
| siCTRL/siALK1 | 1.4 | 0.096 | 0.072 | 3.09 | 0.165 | 1.623 |
| siCTRL/siSMAD4 | 2.92 | 0.143 | 0.131 | 6.45 | 0.246 | 2.955 |

During the first 36 hours of exposure to flow, simulations of control only cells had more vessels with flow dropout than both control/SMAD4 knockdown and control/ALK1 knockdown simulations (Fig. 2A). After 36 hours, the number of vessels with flow dropout in control/SMAD4 knockdown cell simulations increased to greater than control only cells (28.6+/-9.7% and 19.9+/-12.9% respectively at 120 hours in flow). Flow dropout in the control/ALK1 knockdown cell simulations remained lower until 60 hours in flow after which it increased until it was a similar proportion to control/control cells.

**Figure 2.**
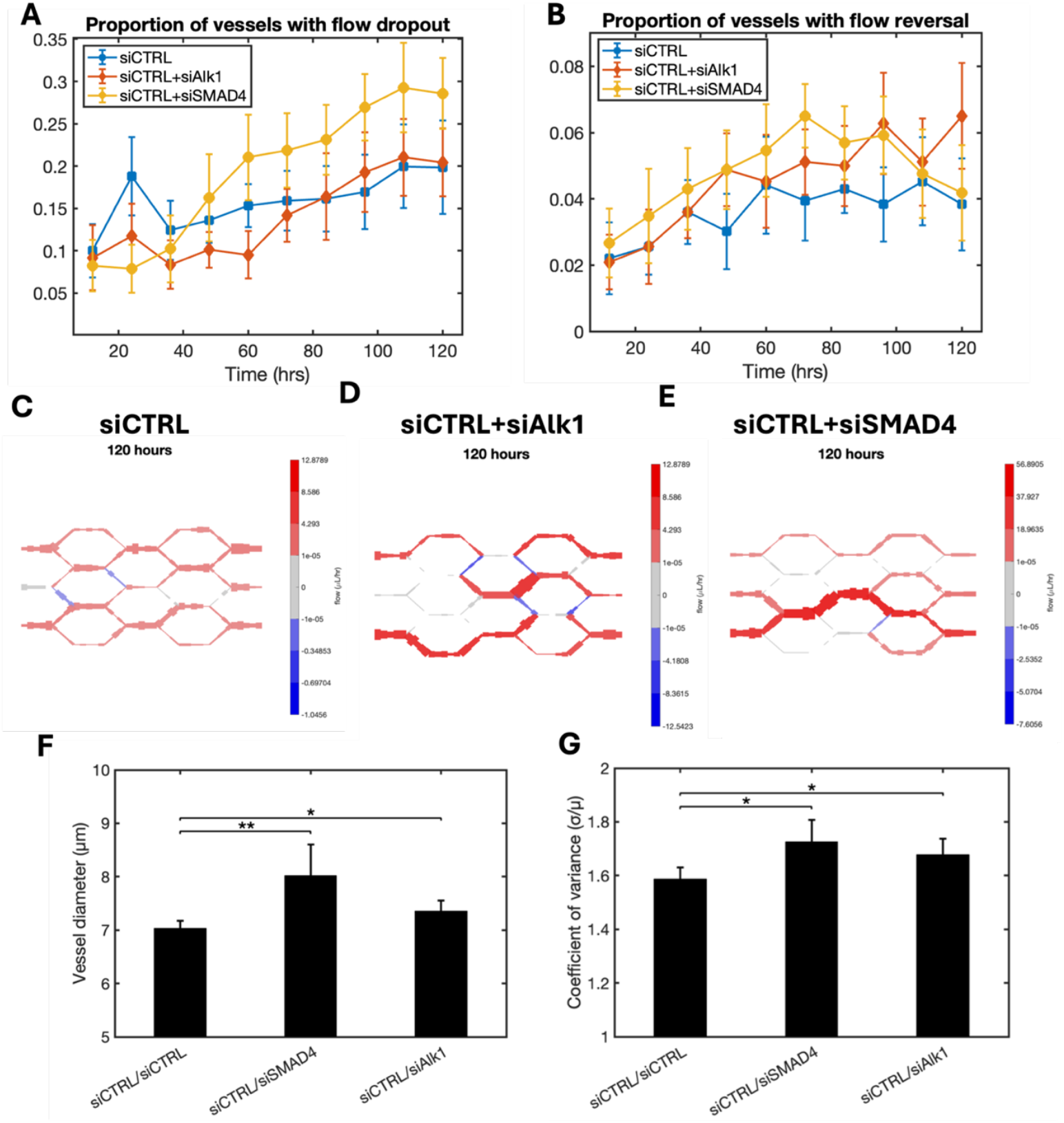
*Figure 2:* Comparison of AVM associated features in honeycomb vessel networks of endothelial cells. (**A**) Proportion of vessel segments with magnitude of flow less than 10^-5^uL/hr. **(B)** Proportion of vessel segments with flow reversal compared to original flow direction (n=21, mean +/-95% CI). (**C-E**) Example images of simulated blood flow magnitude and direction and vessel thickness in vessel networks after 120 hours of development with endothelial cell force coefficients proportional to those inferred from in vitro experiments. (**F**) Mean vessel diameter in perfused vessel segments after 120 hours of development. (**G**) Mean coefficient of variation in perfused vessel segments after 120 hours of development. Plots A, B, F and G show mean +/-95% CI with n = 21.

The proportion of cells with reversal of flow direction compared to the initial state was greater in control/SMAD4 knockdown cell simulations than control simulations over the first 96 hours of simulation (Fig. 2B) after which it decreased to a similar proportion as control cells. Vessels in control/ALK1 knockdown cell simulations initially had a similar proportion of reversed flow to control cells though this proportion increased gradually over time while reversed flow in control cell vessels plateaued leading to a greater proportion of reversed flow in control/ALK1 knockdown cell simulations later on (6.5+/-3.7% in control/ALK1 knockdown and 4.2+/-3.4% in control at 120 hours in flow).

Examples of changes in vessel geometry and flow between control and knockdown simulations after 120 hours are shown in Figures 2 C-E. The average diameter of perfused vessels was found to be significantly larger in both control/SMAD4 knockdown (*p=*0.007) and control/ALK1 (*p=*0.024) simulations with control/SMAD4 producing the largest average increase (Fig 2F). Vessel diameters were also found to be less uniformly distributed in networks with ALK1 or SMAD4 knockdown cells than control cells alone with the coefficient of variation in vessel diameter significantly larger in control/SMAD4 (*p=*0.012) and control/ALK1 (*p=*0.04). Progression of simulated average vessel diameter and coefficient of variation in vessel diameter over time are shown in Supplementary Materials Fig S1A-B.

## Discussion

HHT is characterised by vascular abnormalities in the form of AVMs. Disruption of the BMP 9/10 signalling pathway has been linked to AVM formation in both humans^10^ and mice^11^, but the specific contributions of individual pathway components, and the mechanisms by which they act, remain unclear. Here, we combine *in vitro* mosaic endothelial cell cultures with agent-based simulations of collective cell dynamics to dissect the distinct roles of the receptor ALK1 and the transcription factor SMAD4, both of which have been associated with AVM development.

By calibrating cell–cell interaction parameters in the ABM using experimental observations, we show that both ALK1 and SMAD4 knockdowns influence neighbouring wild-type cells, altering the physical interactions that govern collective EC behaviour. Notably, SMAD4 knockdown cells exhibit enhanced motile force generation, while ALK1 knockdowns reduce the forces that normally allow endothelial cells to separate and rearrange within the layer. These represent two mechanistically distinct modes by which intercellular force balance is perturbed.

Incorporating these altered forces into simulations of vascular remodelling, we predict that both knockdowns increase the likelihood of AVM formation, but via divergent mechanisms. SMAD4 knockdown populations were highly polarised against flow and displayed elevated migration velocities, which, over time, are predicted to cause local cell aggregation and AVM formation. These predictions are supported by our simulation findings, where flow dropout increases significantly in SMAD4-deficient vascular networks over time. The characteristic phenotype of increased diameter of a single vessel in SMAD4 knockdown populations predicted by our simulated vessels matches *in vivo* observations of mouse retinal vasculature upon knockout of SMAD4^12,13^.

In contrast, ALK1 knockdowns displayed less consistent polarity, with a higher proportion of cells migrating against one another. However, the weakened neighbour-separating forces in these cells allowed easier cell passage and prevented vessel blockage, despite the emergence of flow reversals.

These observations align with previous simulation studies^5^ that show imbalances between cell-cell adhesion and displacement forces can disrupt EC rearrangements and promote shunting or vessel occlusion. Importantly, our simulations predict that SMAD4 loss leads to an imbalance between motility and adhesion that drives strongly aligned migration, while ALK1 loss disrupts polarity with subtler mechanical consequences, resulting in divergent vascular phenotypes.

From cell mosaic experiments we found that knockdowns of either gene influenced the polarity of neighbouring wild-type cells, causing faster collective alignment relative to control populations. This suggests that local signalling or mechanical feedback between mutant and wild-type ECs may amplify migratory biases and further destabilise vessel structure.

We acknowledge limitations in our modelling framework. Our agent-based simulations do not currently account for additional regulators of EC dynamics, such as angiogenic sprouting in response to VEGF gradients or shear-stress heterogeneity. The simplified vascular-network geometries also limit topological plasticity, including angiogenesis beyond the original geometry. Moreover, other components of the vascular environment known to influence EC behaviour and vessel stability, such as the basement membrane and mural-cell interactions, are not included in our numerical model. Nonetheless, this reduced complexity enables mechanistic dissection of specific force perturbations and provides a computationally efficient platform for testing candidate hypotheses related to AVM formation.

In summary, our combined experimental and computational results indicate that ALK1 and SMAD4 knockdowns lead to AVM-prone vascular phenotypes via distinct perturbations in endothelial force balance. SMAD4 deficiency drives EC aggregation and vessel obstruction through excessive motility and polarised migration, while ALK1 loss promotes flow reversal through impaired polarity and reduced capacity for neighbour displacement. Together, these findings demonstrate how distinct genetic mutations converge on vascular pathology through disruptions in collective cell dynamics, providing a framework for linking molecular signalling defects to emergent biomechanical behaviours.

## Materials and Methods

### Cell Culture

HUVECs were commercially obtained and cultured in MV2 media (PromoCell). Cells were cultured in gelatinized cell culture flasks and used at passages 2-4. Cells were kept at 37 degrees in a 5% CO_2_ humidity incubator. Knockdown of target genes was achieved by transfecting HUVECs at 60-70% confluence using 10nM siRNA for siCTRL (Qiagen AllStars negative control), siSMAD4 (FlexiTube GeneSolution for SMAD4, Qiagen) or siALK1 (FlexiTube GeneSolution for ALK1, Qiagen) and RNAiMax transfection reagent (Thermofisher) diluted in antibiotic free transfection media (EBM2+OptiMEM) for 5h. Cells were labelled the next day with CellTracker Green or Red (Thermofisher) prior to seeding into flow slides. Briefly, cells were washed once with sterile PBS, and incubated for 45 minutes at 37C with 1:5000 CellTracker diluted in EBM2 media to a final concentration of 2nM. At the end of the incubation, labelling media was aspirated and cells were washed 3 times with MV2 media. Cells were then harvested and mixed at 1:1 ratio, and seeded onto gelatinized 0.4 Luer slides (Ibidi) at a concentration of 2 million cells/ml. Slides were incubated overnight to allow cells to attach and form a monolayer. Further details of the cell culture methodology used are described in Oppenheim *et al*. (2025)^13^ as well as results of qPCR and Western blot assays illustrating the degree to which siRNAs reduce protein expression.

### Shear stress experiments

MV2 growth media was aspirated from slides and replaced with CO_2_ independent media (Promocell basal media without phenol-red and without sodium bicarbonate, supplemented with MV2 supplement kit, B-glycerolphosphate at a final concentration of 4.32ug/ul and sodium bicarbonate at a final concentration of 0.0075%). Slides were connected to fluidic units (Ibidi) and placed within the incubation chamber of the microscope. Shear stress of 0.6 Pa was applied 1 hour after start of imaging for approximately 2.5 days.

### Microscopy

Slides were imaged in pairs using a Zeiss 980 Confocal inverted microscope with a 20x air objective. Imaging setup included 2 selected positions in the middle of every slide; each position was set up with 3x3 tiles and 5-slice Z-stack. Two acquisition green and red channels were set up using the 488 and 561 lasers, respectively. To avoid photo-bleaching, laser intensity was minimized, and signal detection was enhanced by enlarging the pinhole size to 4 AU. Live imaging was defined as a time series of 500 cycles with 7.5-minute intervals. Imaging started one hour prior to shear stress application. CZI files were exported to Fiji. Maximum intensity projection was applied to each position, and files were saved in TIF format.

### Image segmentation

Live images of fluorescent cells were segmented after 36 hours of exposure to unidirectional flow. Cells were segmented using Labkit for ImageJ^8^. Segmented images were manually inspected to add centroids of unlabelled cells and correct mislabelled cells. Segmented cells were tracked over 4 subsequent frames (30 minutes total) using Trackmate for ImageJ. Cell fluorescence colour and centroid position were recorded for each timeframe and distance moved parallel and perpendicular to flow was measured between the first and last frames. Cells were labelled as migrating with or against flow depending upon whether the component of cell displacement parallel to flow was positive or negative.

### Agent-based model

The ABM is described in detail in Edgar *et al*.^5^ ECs are represented as agents consisting of nested ellipses. EC agents migrate in the direction of polarisation at a speed determined by the *coefficient of migration force* (*k*_*mig*_). If neighbouring EC agents are positioned such that their outer ellipses are overlapping, they experience a cohesive force representing cell-cell tension determined by the *coefficient of cohesion* (*k*_*coh*_) and scaled magnitude of the overlap. If EC agents are positioned such that their inner ellipses overlap, they transmit an extrusive force representing cell-cell compression determined by the *coefficient of extrusion* (*k*_*ext*_). Following the initial movement of cells at each timestep, the forces are then recalculated, and cells are intercalated iteratively until all forces are in equilibrium. The new cell positions are given as an output.

Since the effect of cell forces on vessel dynamics is dependent upon the relationship between extrusion or cohesion forces and migration force, the force coefficients can be combined as ratios to represent the work required to elicit cell behaviours. The work required to push two adjacent cells together is given by:

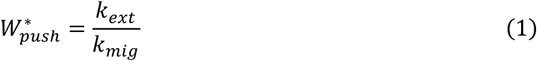

While the work required to pull two adjacent cells apart is given by:

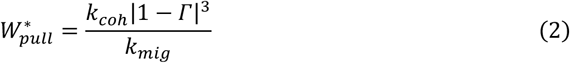

Where Γ is the yield stretch, the maximum stretch ratio of a cell before junction connections are lost. Γ was set to a constant value of 1.5 throughout. The ABM contains a ‘flow sensitivity’ parameter regulating the time taken for cells to become polarised with flow. In this study, it was also assumed that after 36 hours in flow all cells would begin polarised in parallel with the direction of flow.

For inference of intercellular force coefficients from *in vitro* cell experiment, the geometrical constraints of the ABM were set to a flat plate with dimensions dependent upon the size of the experimental image for which they are simulating. Cells were seeded in simulations depending on the location of their centroids measured from the corresponding *in vitro* experiment. Cell length was taken from an average of measurements of segmented *in vitro* cells. Cells leaving one edge of the geometry enter on the opposite edge.

For simulations of vascular networks, cells were seeded into an idealised ‘honeycomb’ plexus of bifurcating vessels^5^. Individual vessels were formed from a 50 μm length with 5 discrete cylindrical segments and each segment initiated with 4 ECs of length 10 μm and width 5 μm. Vessel radius changes depending upon the flux of cells into and out of a lumen segment. Pressurised fluid is simulated within the vessel lumens, with pressure stored at nodes between luminal segments. Flow at inlets and outlets is at constant pressure but changes dynamically within the network with changes in vessel radius. A flow magnitude of less than 10^-5^ μl/hr within a vessel is defined as “no flow” and recorded as flow drop out.

### Approximate Bayesian Inference

Values of *k*_*mig*_, *k*_*coh*_ and *k*_*ext*_ are inferred from experimental measurements of cell displacement using approximate Bayesian computation with a sequential Monte Carlo sampling method (ABC–SMC)^9^. The ABM is run with input parameter values sampled from a prior distribution within a reasonable range (we apply uniform distributions with range 0-1 for *k*_*coh*_ and *k*_*ext*_ and 0-5 for *k*_*mig*_). The vertical (perpendicular to flow) and horizontal (parallel to flow) distances between simulated cell locations as outputs from the ABM and those measured from *in vitro* experiments is calculated and used to generate a loss function to determine the accuracy of that simulation. ABC–SMC runs multiple simulations with varying input parameter values and selects samples from which the loss function is below a threshold (ε) to build a probability distribution for likely parameter values. ε is systematically decreased, and so, if a solution is possible, the probability distribution will converge towards the most likely values for each of the parameters.

The loss function for comparison of experimental and simulated cell displacement is as follows:

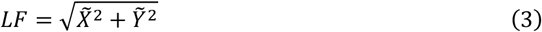

Where

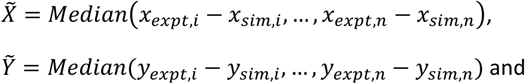

*x*_*expt,i*_, *y*_*expt,i*_ are experimentally measured centroid coordinates of cell *i* in the directions parallel and perpendicular to flow respectively and *x*_*sim,i*_· *y*_*sim,i*_ are simulated centroid coordinates of cell *i* in the directions parallel and perpendicular to flow respectively.

### Application of flat plate force coefficients to an idealised vascular plexus

Once *k*_*mig*,_ *k*_*coh*_ and *k*_*ext*_ have been inferred from *in vitro* experiments on flat plates, the relative differences in their values are applied as parameter inputs to the ABM with the simplified ‘honeycomb’ vascular network geometry. In previous simulations of ECs in a vascular plexus^5^, intercellular force coefficient values for *k*_*mig*_=3, *k*_*coh*_ =1/3 and *k*_*ext*_ =3 were found to generate stable networks, free from adverse vascular remodelling and with cell migration speeds similar to those found *in vivo*. Intercellular force coefficients for control cells were therefore set to these values. Values for intercellular force coefficients in vessel networks with control and knockdown cells were calculated using the following equation:

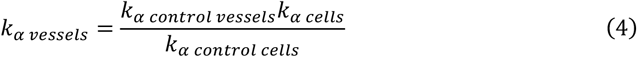

Where *k*_*α_control_vessels*_ and *k*_*α*_*vessels*_ are force coefficients for vascular plexus simulations for control and control/knockdown experiments respectively and *k*_*α*_*control_cells*_ and *k*_*α_cells*_ are force coefficients inferred from *in vitro* experiments for control and control/knockdown experiments respectively.

Edgar *et al*.^5^ also found that the formation of shunts in simulated vessel networks analogous to AVM was associated with loss of perfusion and/or reversal of flow in vessels and so measurement of perfusion loss and flow reversal in vessels are used here as indicators of AVM formation. To assess AVM-like vascular remodelling, the mean of all vessel diameters in perfused vessels was recorded as well as the coefficient of variation of vessel diameters (standard deviation in vessel diameter/mean vessel diameter), an indicator of uniformity of vessel width.

A set of 21 random seed numbers were generated and used to randomise the starting positions for cells within the simulated vascular plexus. Each starting position was used to simulate cells with cell force coefficients and cell direction of motion matching control, ALK1 knockdown and SMAD4 knockdown ECs.

## Statistics and code availability

Results for measured and inferred values from *in vitro* cell experiments are shown +/-1 standard deviation from the mean. Features of ABM vessel network simulations are shown +/-95% confidence interval. Two-sample t-tests were used to assess significance of differences in vessel. Statistical analysis was conducted in Matlab.

The agent-based modelling code, together with scripts linking the ABM to pyABC and example tracking datasets, is available at https://github.com/dhardma2/EC_ABM

## Acknowledgments

We thank Lowell Edgar for developing the original version of the agent-based modelling scripts used and adapted in this study.

