## Supplemental Table S1, Supplemental Figure S1 for "Disrupted endothelial cell–cell dynamics in ALK1 and SMAD4 deficiency drive arteriovenous malformations"

### Supplementary materials

| Experiment | Cell type | Mean percentage of cells moving in direction of flow (+/- 1SD) | | |
|---|---|---|---|---|
| | | **36 hrs in flow** | **48 hrs in flow** | **60 hrs in flow** |
| **siCTRL/siCTRL** | siCTRL cells (red) | 40.8 +/- 5.3 | 31.1 +/- 2.6 | 21.8 +/- 3.7 |
| | siCTRL cells (green) | 39.1 +/- 2.4 | 32.2 +/- 3.0 | 25.8 +/- 5.4 |
| | **Combined** | **40.0 +/- 4.3** | **30.9 +/- 1.1** | **23.7 +/- 4.6** |
| **siCTRL/siALK1** | siALK1 cells | 33.4 +/- 5.2 | 22.5 +/- 6.5 | 23.6 +/- 6.9 |
| | siCTRL cells | 35.4 +/- 5.4 | 27.9 +/- 5.9 | 23.3 +/- 4.9 |
| | **Combined** | **34.4 +/- 5.4** | **24.9 +/- 6.1** | **23.6 +/- 5.8** |
| **siCTRL/siSMAD4** | siSMAD4 cells | 14.4 +/- 2.8 | 14.5 +/- 3.5 | 17.1 +/- 3.6 |
| | siCTRL cells | 22.1 +/- 5.2 | 20.1 +/- 4.2 | 22.8 +/- 6.0 |
| | **Combined** | **17.4 +/- 2.4** | **16.8 +/- 1.2** | **19.5 +/- 4.1** |

*Table S1: Proportional cell direction of motion parallel to flow over time.*

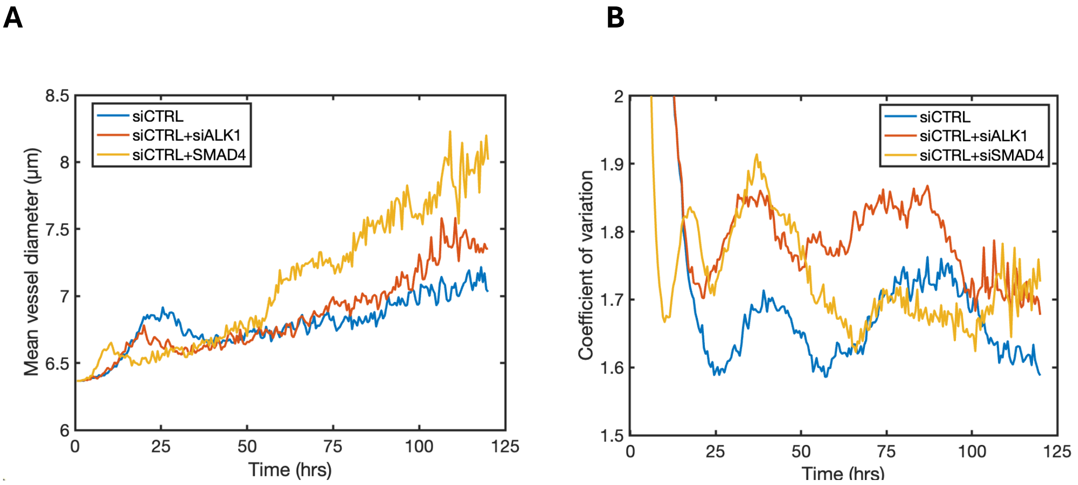


*Figure S1: Changes in average vessel geometry over time. (A) Mean vessel diameter in perfused vessel segments. (B) Mean coefficient of variation of vessel diameter, an indicator of uniformity of distribution.*
